# Using Mixture Density Networks to Predict Plot-level Environmental Factors from Tree Secondary Metabolites

**DOI:** 10.64898/2026.11.24.768708

**Authors:** Kelly To, Dale L. Forrister, Brian E. Sedio, Claus O. Wilke

## Abstract

Tropical forests contain remarkable phytochemical diversity, but the relationship between diverse plant metabolites and underlying climatic and environmental variation remains poorly understood. Mixture density networks (MDNs) offer a flexible approach for predicting variables with multimodal distributions. Traditional regression models produce single-value predictions, which may fail to capture the full variability of widely distributed variables. We evaluated the performance of an MDN model on a Panama metabolomics dataset, in which each metabolite occurs across multiple forest plots, resulting in environmental variables with multiple associated values. The MDN recovered the overall range and broad shape of environmental distributions, although feature-specific predictions often converged toward shared distributional patterns. These results suggest that mixture density networks can capture coarse environmental structure from metabolomic representations, while also highlighting challenges associated with limited environmental variation and sparse observations.

## Introduction

Lowland tropical forests are among the most biologically diverse ecosystems on Earth, often containing more plant species in a single few hectares than can be observed in all of temperate North America (Valencia et al., 2004). This diversity extends beyond plant species to include different aspects of plant functional diversity, most noticeably phytochemical diversity. Individual plants produce hundreds to thousands of unique secondary metabolites (Forrister et al., 2023), and collectively the plant kingdom is estimated to produce more than a million unique secondary metabolites (Afendi et al., 2012). This chemical diversity is believed to be highest in tropical forests, in particular, where plants invest in both high diversity and abundance of secondary plant metabolites. Plant secondary metabolites play critical roles in modulating ecological interactions, from pollination to attack by insect herbivores and fungal pathogens (Coley et al., 1996; Endara et al., 2023). As a result, individual metabolites may be associated with a wide range of environmental contexts, producing distributions of environmental conditions that are often broad or multimodal.

When environmental conditions associated with individual metabolites exhibit broad or multimodal distributions, mixture density networks (MDNs) offer an alternative to conventional regression models by predicting an entire probability distribution rather than a single point estimate. MDNs model multimodal relationships by assuming that the target distribution can be represented as a mixture of normal distributions, with the network predicting the parameters of each component (Bishop, 1994). Although MDNs remain uncommon in ecological modeling, they have been applied in other fields, such as electrical load forecasting and wind speed/power prediction (Vossen et al., 2018; Men et al., 2016).

In this study, we explore whether MDNs can learn ecologically meaningful relationships between plant secondary metabolite composition and environmental conditions in a tropical forest. We analyze an untargeted metabolomics dataset derived from 455 species of tropical trees which are distributed across a network of 44 forest plots in central Panama (Condit, Engelbrecht, Pino, Pérez, et al., 2013), a region characterized by strong edaphic and precipitation gradients that occur over short distances across the narrow Isthmus. Due to these gradients, Panama’s forests harbor some of the highest levels of species *β*-diversity in the tropics (Condit, Pitman, et al., 2002). Using DreaMS-derived (Bushuiev et al., 2026) embeddings of MS/MS spectra, we ask whether metabolite chemistry can be used to recover distributions of plot-level environmental conditions, including dry-season moisture and soil phosphorus metrics. Because individual metabolites occur across multiple plots, associated environmental values form distributions that are often wide and multimodal, making this data set particularly suitable for evaluating a mixture density network.

We evaluate the performance of our MDN on both common and rare metabolites that occur in only a few plots, allowing us to assess how data availability influences distributional prediction. Our results show that MDNs can recover key aspects of the observed environmental distributions and, in many cases, capture feature-level variation even under data-limited conditions. Together, these findings highlight both the potential and the limitations of mixture density networks for linking metabolomic data with environmental gradients in ecological systems.

## Methods

### Forest plots and environmental variables

Our data consists of MS/MS spectra and plot-level environmental metadata from tree leaf samples collected across 44 forest plots in central Panama (Figure 1a). This region is notable for its steep gradient in plant-available phosphorus, showing more variation than what has been measured across the Amazon basin. This gradient is ecologically important, as dry-season moisture and soil phosphorus have been identified as key predictors of tree distributions in the region (Condit, Engelbrecht, Pino, Pérez, et al., 2013).

**Figure 1:**
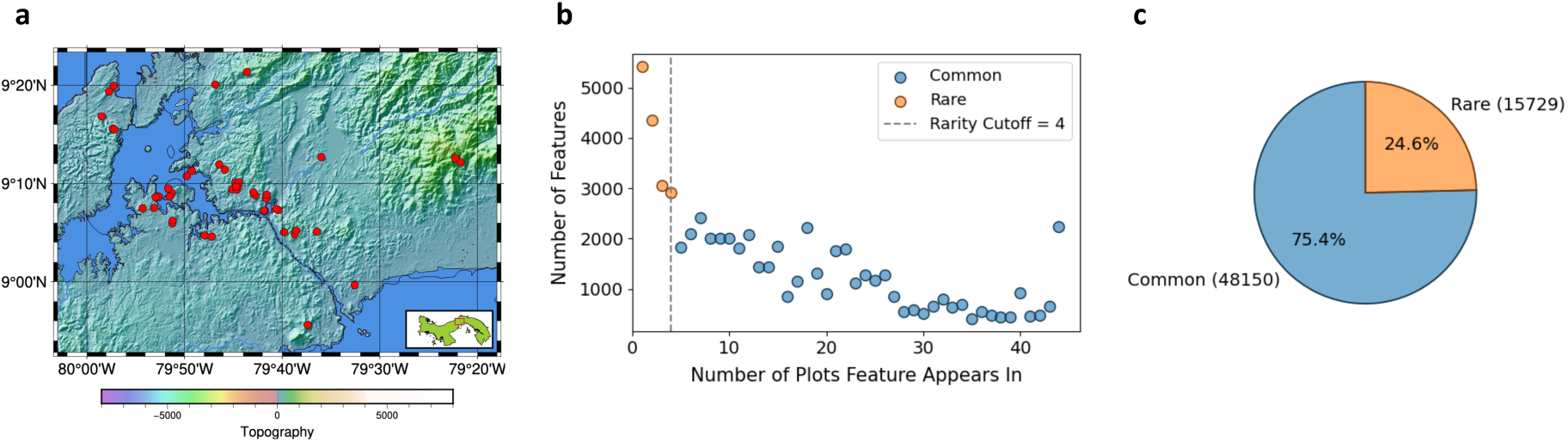
(a) Map of the study region in Panama. Locations of the leaf collection plots are indicated by the red dots. Most features appear in multiple plots. (b) Number of features repeated across multiple plots. (c) Relative breakdown of rare and common features. Rare features are defined as occurring in at most 4 distinct plots.

Based on these findings, we focused on three plot-level environmental variables: dryseason moisture, resin phosphorus, and total phosphorus. Dry-season moisture captures the amount of water lost from the soil during periods when evapotranspiration exceeds precipitation. Resin phosphorus reflects the amount of plant-available phosphorous in the soil, while total phosphorus represents the overall phosphorus content in the soil (Condit, Engelbrecht, Pino, Pérez, et al., 2013).

Tree species abundances were obtained from the Panama forest plot network (Condit, Engelbrecht, Pino, Pérez, et al., 2013; Condit, Pérez, et al., 2022), part of the Forest Global Earth Observatory (Davies et al., 2021). This network comprises 65 plots near the Panama Canal established between 1994 and 2014, most of them 1 ha in area (range 0.32–6 ha). In each plot, all free-standing woody stems *≥* 1 cm diameter at breast height (1.3 m) were mapped, measured, and identified to species. For each plot, we used the most recent census and retained only stems recorded as alive. Environmental data for the plots came from Condit, Engelbrecht, Pino, Pérez, et al. (2013) and Condit, Engelbrecht, Pino, Turner, et al. (2013), which measured total phosphorus and biologically available resin phosphorus in soil samples collected in each plot. They also used records from 47 rainfall gauges maintained by the Panama Canal Authority to calculate the maximum annual cumulative moisture deficit (precipitation minus potential evapotranspiration), and then interpolated these values with a spatial kernel to estimate the dry-season moisture deficit at each plot. We restricted our analyses to the 44 plots for which both soil phosphorus and moisture deficit estimates were available.

### Sampling

We collected expanding, unlignified leaves that were between 50 and 90% fully expanded from saplings, shrubs, and understory trees between March 2018 and December 2021. Three to five leaves from each individual were placed in a manila coin envelope and immediately flash frozen in liquid nitrogen in the field using a portable dewar (PrincetonCryo, Pipersville, PA, USA) and freeze-dried upon return to the laboratory. We sampled 5 individuals per species for a total of 413 species recorded in the forest plots. Freeze-dried leaf samples were shipped to the University of Texas at Austin for extraction and analysis using untargeted metabolomics.

### Untargeted metabolomics

Chemical extraction and analytical chemistry instrumental methods followed Sedio et al. Sedio et al. (2021). Briefly, a mass of 50 mg of dried leaf tissue from each sample was pulverized and homogenized using a Qiagen TissueLyser ball mill (Hilden, Germany). From this, a 10-mg subsample was weighed (*±*0.01 mg) and extracted in 1800 *µ*l 90:10 methanol:water pH 5 (vol:vol) overnight at 4 *^◦^*C. Extracts of up to 5 individuals per species were pooled for analysis.

Species pools were filtered and analyzed using ultra-high performance liquid chromatography-heated electrospray ionization-tandem mass spectrometry (UHPLC-HESI-MS/MS) using a Thermo (Waltham, MA, USA) Vanquish UHPLC and QExactive quadrupole-orbitrap high-resolution MS. Metabolites were separated for 20 min using a C-18 column and ionized in positive mode using full-scan MS1 and data-dependent acquisition of MS2 spectra (Sedio et al., 2021). Spectra were deposited as a public MassIVE dataset on the Global Natural Products Social (GNPS) Molecular Networking server (Dataset MSV000104086).

Raw MS spectra were centroided and peaks detected and aligned using MZmine (Schmid et al., 2023). Aligned chromatograms were used to generate a feature-based molecular network (Nothias et al., 2020) using GNPS (Wang et al., 2016) to generate a consensus spectrum for each unique detected metabolite signal, exported as an .mgf file. These signals are commonly referred to as “features,” meaning that in this study, a feature represents a unique MS/MS spectrum corresponding to a metabolite detected across one or more forest plots. Since the same metabolite can occur in multiple plots, each feature may be associated with multiple measurements of a given environmental variable, producing a distribution of values for a single metabolite/feature.

To characterize how broadly each feature was distributed, we examined the number of plots in which each feature appeared. The distribution shows a steep initial decline that transitions to a more gradual decrease after about four plots, followed by an uptick in features present in all 44 plots (Figure 1b). Based on this pattern, we constructed two training sets, one derived from all features and another from only rare features. Rare features were defined as those found in at most four of the 44 plots, a cutoff chosen to align with the declining trend of the distribution and to evaluate model performance on infrequently observed features. The final dataset consisted of 63,879 features. Most of the features appeared in more than four plots, with roughly three-quarters (48,150) classified as common and the remaining quarter (15,729) classified as rare (Figure 1c).

### DreaMS embedding generation

We generated embeddings for the MS/MS spectra using DreaMS, a neural network designed for interpreting MS/MS data, following the procedures outlined in the documentation available at: https://dreams-docs.readthedocs.io/en/latest/. DreaMS produced a 1024-dimensional embedding for each feature and stored the results in a single .hdf5 file (Bushuiev et al., 2026).

### Data preprocessing

Prior to model training, we constructed a unified feature-level dataset by merging the data sources from above. The merges were performed using the shared FEATURE ID key for metabolite features and plot identifiers for environmental variables. First, we merged the DreaMS embeddings with the feature-by-species matrix and reshaped the dataset into long format, retaining only nonzero feature abundances. We repeated this process with the feature-by-plot matrix, producing a long-format table of feature abundances across plots. This table was then merged with the plot metadata to attach environmental variables to each observation. The resulting combined dataset contained, for every observed metabolite–plot occurrence, its DreaMS embedding, abundance, and the associated environmental measurements.

We then selected our environmental target variables and filtered the dataset to retain only the relevant columns. Each row corresponded to a single feature occurrence, containing the FEATURE ID, DreaMS embedding, and environmental variables measured at the plot where the feature was observed. For each target variable, we constructed two datasets: one containing all features and another restricted to rare features. Rare features were defined as metabolites observed in four or fewer plots; these were isolated by grouping the full dataset by FEATURE ID and filtering accordingly. Both datasets were then split into training, validation, and test subsets using an 80/10/10 grouped split by FEATURE ID, ensuring that all observations of a given feature were assigned to the same subset. The resin phosphorus and total phosphorus data for rare features were log transformed using np.log(). Each subset was stored as a dictionary containing PyTorch tensors for feature IDs, embeddings, and target variables, and all subsets were saved together in a single .pt file for use during model training.

### Model architecture

The model we used is a mixture density network, in which a feed-forward neural network outputs the parameters of a Gaussian mixture model. The mixture model represents the conditional probability density *p*(*y|x*) of the target variables given an input vector *x* as a weighted sum of *K* Gaussian components (Bishop, 1994). For each component *k* = 1*, . . ., K*, the MDN predicts the mixture weight *π_k_*(*x*), mean *µ_k_*(*x*), and variance *σ_k_*(*x*).

MDNs are trained by minimizing the negative log-likelihood of the mixture distribution (Guillaumes, 2017):

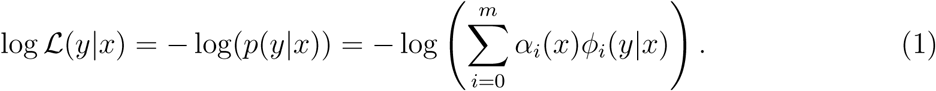

Our MDN implementation was adapted from a PyTorch version available at: https://github.com/tonyduan/mixture-density-network/tree/master. To improve numerical stability, we modified the computation of (*σ*) by moving the small constant after exponentiation. This ensures that *σ* is positive, preventing near-zero variance estimates that could cause unstable gradients or NaN losses during training.

### Model training

For each target variable, we trained two versions of the MDN: one using all features and one using only the rare features. In both cases, we used a batch size of 32 and trained for up to 100 epochs with early stopping (patience of 20 epochs). For the model architecture parameters, we used a hidden dimension of 64 and selected the number of Gaussian mixture components through empirical evaluation. We compared rare-feature models trained with one to five, 10, 15, and 20 components, analyzing their training and validation losses across the three target variables. Through this, we decided on the configuration with 15 components for a balance between negative log-likelihood and distributional smoothness, as additional components improved loss but produced increasingly peaked distributions that risked overfitting. For consistency, the same number was used for the all-features model. Training used a learning rate of 0.0004, a weight decay of 1 *×* 10*^−^*^6^, Adam optimizer betas of 0.9 and 0.95, and a dropout rate of 0.1. The best three checkpoints were saved.

### Prediction generation

After training, we generated predictions for each target variable using the best-performing checkpoint of each model. Predictions were obtained by sampling from the MDN’s learned mixture distribution via the samples() function. To assess the predictions of individual features, we drew 100,000 samples per feature, providing a high-resolution approximation of each feature’s predicted conditional distribution. These samples were used to construct predicted density curves, which were then compared against the corresponding observed target variable values.

## Results

### Modeling approach

To link metabolite chemistry with environmental variation, we combined DreaMS embed-dings with plot-level environmental metadata and trained a mixture density network to model their relationship (Figure 2a). MS/MS spectra were first processed using DreaMS, a neural network that interprets spectral data (Bushuiev et al., 2026). Each metabolite was represented as a 1024-dimensional embedding that captures its spectral information. These embeddings were then paired with the selected plot-level environmental variables and used as input to the MDN, which models the conditional distribution of each target variable given a metabolite’s embedding.

**Figure 2:**
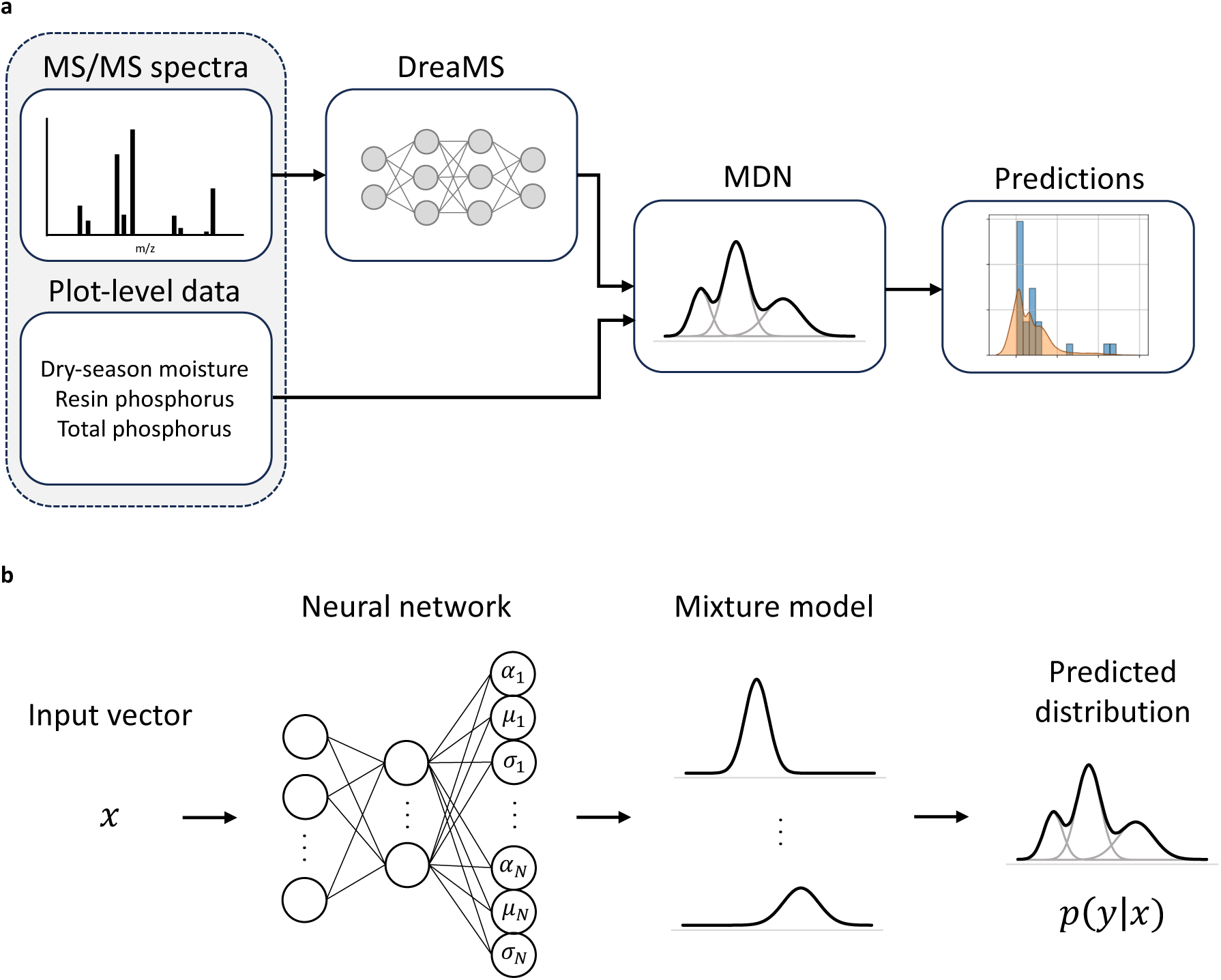
(a) Schematic overview of the modeling approach. MS/MS spectra and plotlevel environmental metadata were collected from leaf samples. Spectra were processed with DreaMS to generate embeddings for each metabolite. Embeddings were paired with environmental metadata and used to train a mixture density network. The trained model predicts environmental variables for each metabolite. (b) A mixture density network consists of a feed-forward neural network followed by a Gaussian mixture model. Given an input vector *x*, the neural network outputs the mixture weight *π_k_*(*x*), mean *µ_k_*(*x*), and variance *σ_k_*(*x*) of each mixture component *k*. These parameters define the individual component density curves in the mixture model. Summing the weighted components yields the overall conditional density *p*(*y|x*), which represents the full predicted distribution of the target variable.

A mixture density network consists of a feed-forward neural network that outputs the parameters of a Gaussian mixture model (Bishop, 1994). For each mixture component *k*, the network predicts a mixture weight *π_k_*(*x*), mean *µ_k_*(*x*), and variance *σ_k_*(*x*), allowing the model to capture complex and potentially multimodal relationships between metabolites and environmental variables (Figure 2b). Rather than outputting a single predicted value, the trained MDN provides the full conditional probability distribution *p*(*y|x*). This allows the model to represent uncertainty, multimodality, and non-linear associations between metabolites and environmental variables.

### Model performance on all features

We first trained the model using all features to evaluate its performance across metabolites occurring in any number of plots. We chose to log transform the data for resin phosphorus and total phosphorus to make the actual distributions more normal. Because MDNs are trained by minimizing the negative log-likelihood, more negative loss values indicate better model fit (Guillaumes, 2017). Training and validation losses decreased for all three target variables relative to epoch 0. However, the reductions were modest, suggesting that the model may have had some difficulty learning an effective representation (Figure 3). The best-performing checkpoints occurred at epoch 13 for dry-season moisture, epoch 2 for log resin phosphorus, and epoch 2 for log total phosphorus.

**Figure 3:**
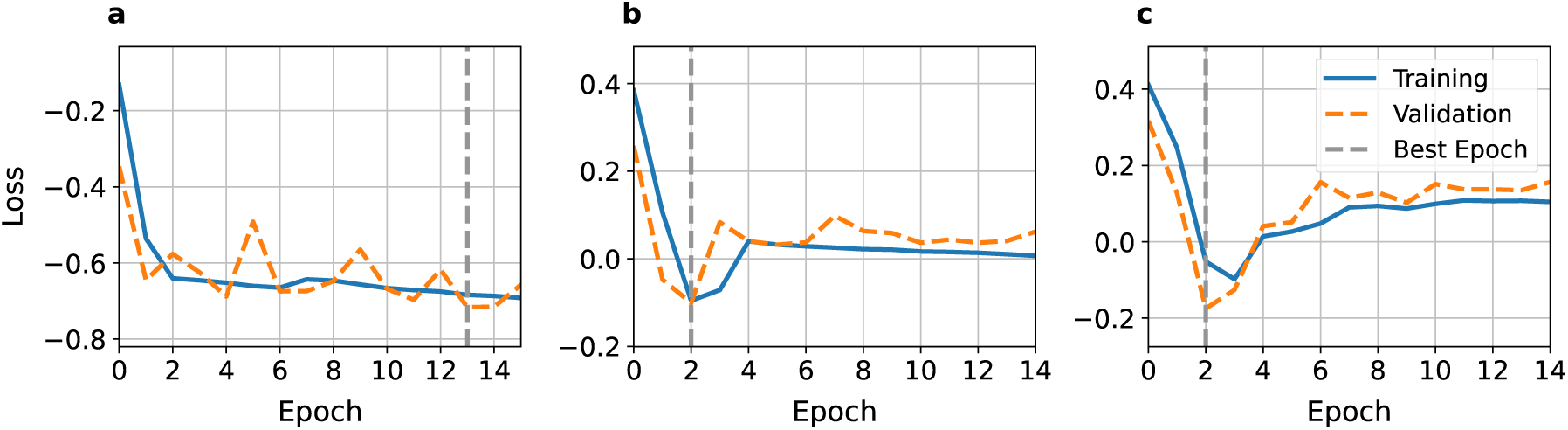
Training and validation loss curves for the 3 target variables when training on all features. The blue solid lines show the training loss, while the orange dashed lines show the validation loss. The gray dashed vertical lines indicate the epochs with the lowest validation loss. (a) Loss curves for dry-season moisture. (b) Loss curves for log resin phosphorus. (c) Loss curves for log total phosphorus.

To assess predictive performance, we generated predictions for every sample in the test set and compared the aggregate predicted distributions to the corresponding observed distributions. The predicted distributions captured the overall range and general shape of the observed values for all three target variables (Figure 4).

**Figure 4:**
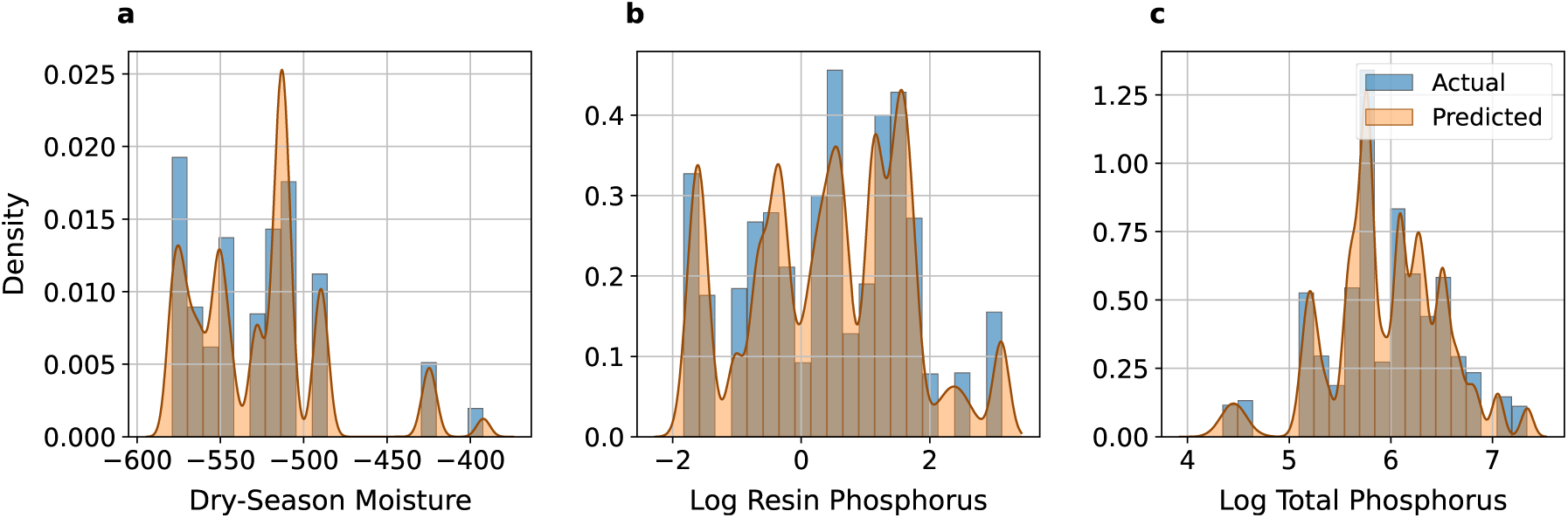
The overall predicted distributions agree well with the overall distributions across features when training on all features. The blue bars show the actual values from the test set for each target variable. The orange curves show the distributions of the model’s predictions. A prediction was generated for every observation in the test set to construct each distribution curve. (a) Actual and predicted distributions for dry-season moisture. (b) Actual and predicted distributions for log resin phosphorus. (c) Actual and predicted distributions for log total phosphorus.

Furthermore, we examined feature-specific predictions to assess how well the model captured variation at the level of individual metabolites. For each target variable, we randomly selected three features from the test set and generated 100,000 samples per feature to estimate the predicted distributions. This large number of samples was chosen to obtain smooth, stable estimates that closely approximated the distributions learned by the models (Supplementary Figure S1). In several cases, the predicted curves were broadly consistent with the observed values for a given feature. However, many predicted distributions exhibited highly similar shapes across different features, even when the underlying feature-level distributions differed (Figure 5, as well as Supplementary Files S2, S3, S4 for more features). This pattern suggests that when training on all features, the MDN may have learned a common distributional structure that is applied across features, rather than capturing feature-level differences.

**Figure 5:**
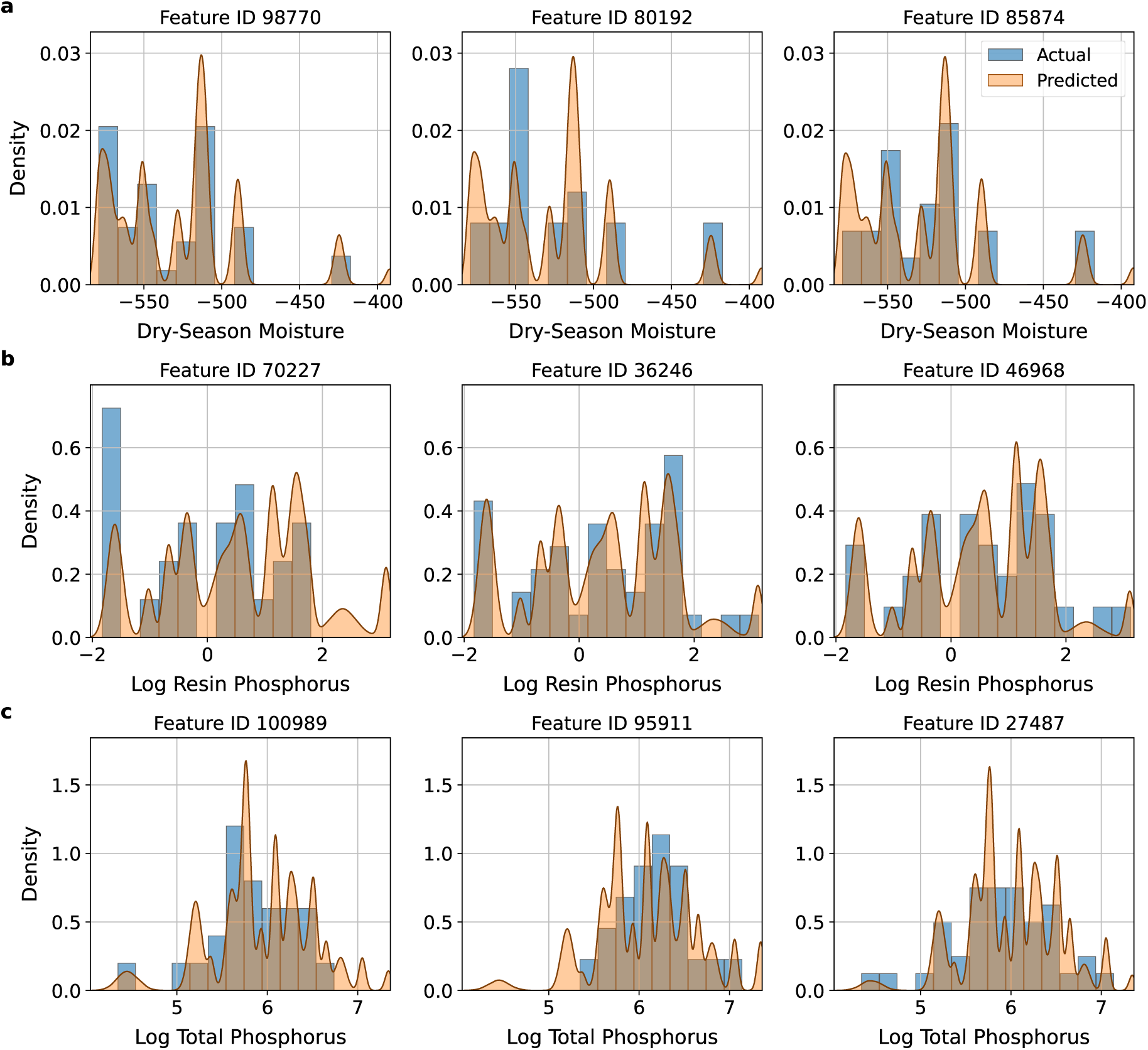
Feature-level predicted distributions align with observed values, with some consistency across features. The blue bars show the actual values of a feature randomly selected from the test set. The orange curves, constructed by sampling 100,000 predictions per feature, show the distributions of the model’s predictions. (a) Actual and predicted distributions of 3 random features for dry-season moisture. (b) Actual and predicted distributions of 3 random features for log resin phosphorus. (c) Actual and predicted distributions of 3 random features for log total phosphorus. For additional examples, see Supplementary Files S2, S3, S4.

### Model performance on rare features

Next, we evaluated the MDN’s performance on rare features that appeared in at most four plots by training the model using only such features. As in the model trained on all features, both training and validation losses decreased across all three target variables compared to epoch 0, reflecting an improved fit to the observed environmental distributions over the course of training. All of the losses were lower than those observed in the all-features model, with log resin phosphorus exhibiting the largest reduction in validation loss, suggesting the greatest improvement in fit to unseen data (Figure 6). The best-performing checkpoints occurred at epoch 28 for dry-season moisture, epoch 36 for log resin phosphorus, and epoch 12 for log total phosphorus.

**Figure 6:**
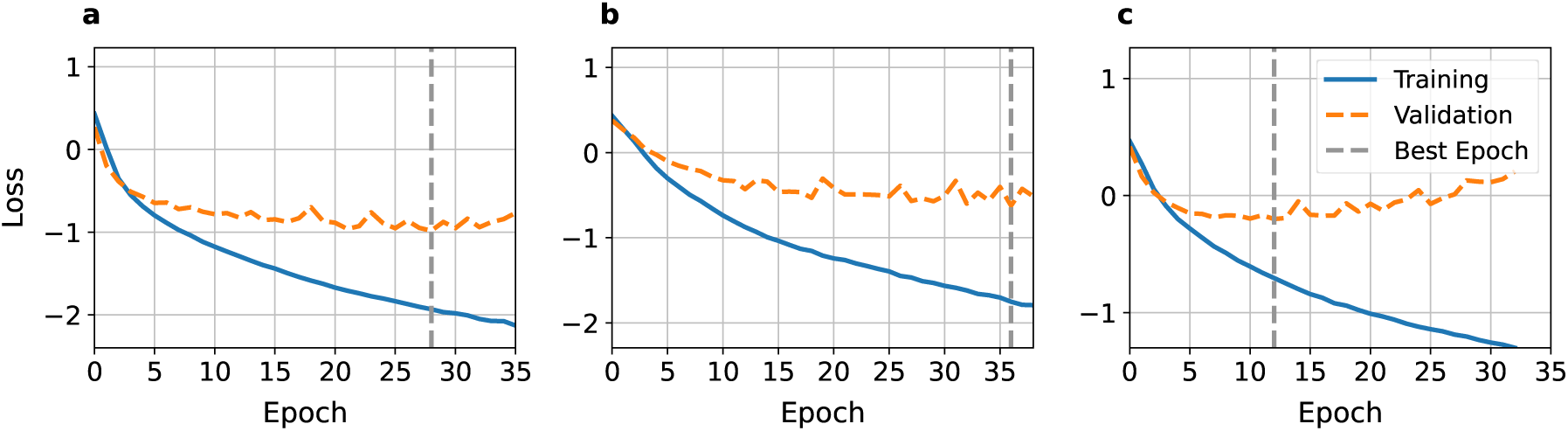
Training and validation loss curves for the 3 target variables after training on rare features. The blue solid lines show the training loss, while the orange dashed lines show the validation loss. The gray dashed vertical lines indicate the epochs with the lowest validation loss. (a) Loss curves for dry-season moisture. (b) Loss curves for log resin phosphorus. (c) Loss curves for log total phosphorus.

Despite the limited number of observations per metabolite, the MDN generated predicted distributions that captured the broad structure of the data. However, it did struggle to reproduce many of the peaks as pronounced as they are in the actual distributions (Figure 7). This indicates that, under data-limited conditions, the model can somewhat recover meaningful distributional structure for the environmental variables.

**Figure 7:**
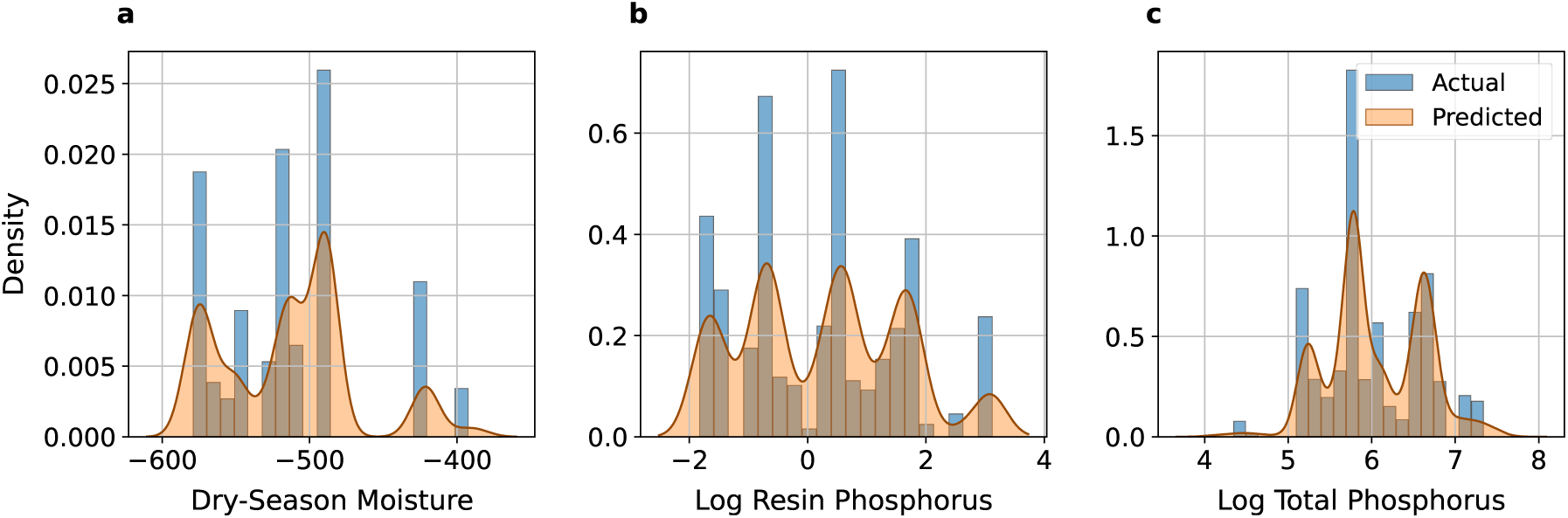
Predicted distributions align more closely with some environmental variables than others when training on rare features. The blue bars show the actual values from the test set for each target variable. The orange curves show the distributions of the model’s predictions. Predicted distributions were constructed by generating one prediction per test observation. (a) Actual and predicted distributions for dry-season moisture. (b) Actual and predicted distributions for log resin phosphorus. (c) Actual and predicted distributions for log total phosphorus.

We also examined feature-specific predictions for the model trained on rare features. For each target variable, we constructed actual versus predicted distribution plots for all features that occurred in exactly four plots. These features were sorted by their mean observed values, from lowest to highest. To illustrate model behavior across this range, we selected three features representing the lower, middle, and upper portions of the distribution. While the model was able to approximate the distribution of some individual features, performance was more variable than for the model trained on all features, with greater discrepancies between predicted and actual distributions for several metabolites (Figure 8, see Supplementary Files S5, S6, S7 for the full sets of features). Oftentimes, the actual values were within the predicted distribution, but the peaks of the distribution did not necessarily represent the actual values well.

**Figure 8:**
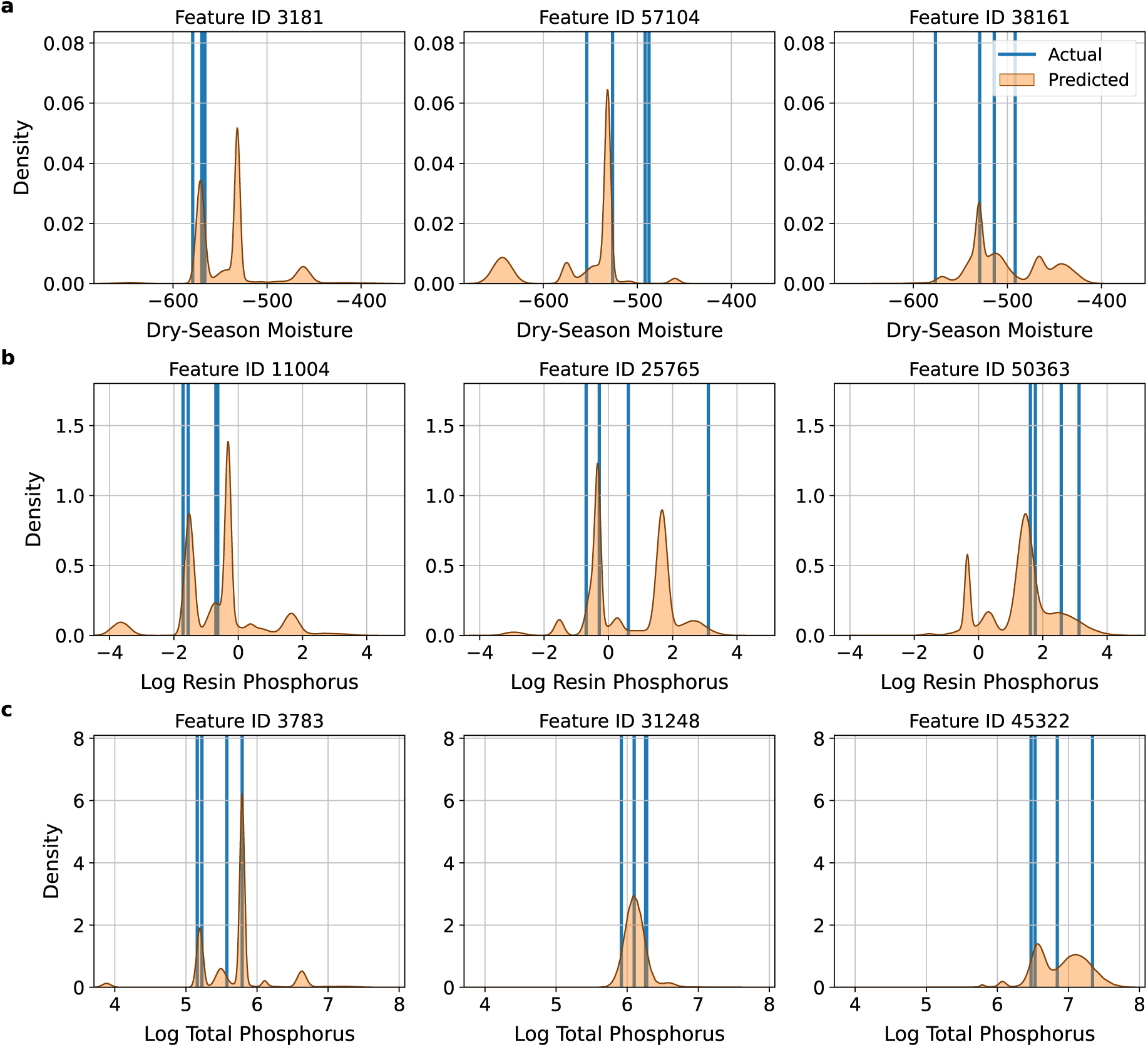
Feature-specific predicted distributions for rare metabolites show variable alignment across the observed range. The blue bars show the actual values of a feature selected from the test set. The orange curves, constructed by sampling 10,000 predictions per feature, show the distributions of the model’s predictions. For each target variable, three features were selected after sorting all rare features by their mean observed values: one from the lower end, one from the middle, and one from the upper end of the distribution. (a) Actual and predicted distributions of 3 features for dry-season moisture. (b) Actual and predicted distributions of 3 features for log resin phosphorus. (c) Actual and predicted distributions of 3 features for log total phosphorus. For the full set of features, view Supplementary Files S5, S6, S7.

## Discussion

In this study, we investigated whether mixture density networks can learn relationships between plant secondary metabolites and plot-level environmental conditions in the tropical forests of central Panama. Using untargeted metabolomics data from tree leaf samples collected across 44 forest plots, we generated DreaMS embeddings from MS/MS spectra and used these embeddings to predict distributions of dry-season moisture, log resin phosphorus, and log total phosphorus. Because individual metabolites can occur across multiple plots, each metabolite may be associated with multiple measurements of an environmental variable. This results in distributions that are often broad and/or multimodal, which are difficult to model with traditional regression approaches. By modeling these as conditional probability distributions rather than single-point estimates, MDNs were able to recover broad distributional patterns present in both the complete dataset and the rare-metabolite subset and better capture the multimodality inherent in the system.

Across all features, the MDN was able to capture the overall distributions of dry-season moisture, resin phosphorus, and total phosphorus reasonably well. The aggregate predicted distributions closely matched observed ranges and general shapes, indicating that the model learned broad environmental structure from the metabolite embeddings. However, feature-level analyses revealed that the predicted distributions often exhibited similar shapes across different metabolites, suggesting that the model partially relied on global distributional cues rather than fully metabolite-specific information.

Training on rare features highlighted both the strengths and limitations of the MDN under data-limited conditions. Even with each metabolite observed in at most four plots, the model was still able to recover some meaningful distributional structure for the environmental variables. Performance was nonetheless more variable across features, with predictions for dry-season moisture being less aligned with observed values. These differences likely reflect both reduced data availability and intrinsic properties of the target variables. Dry-season moisture may integrate complex climatic and edaphic processes that are less directly encoded in secondary metabolite profiles than soil phosphorus metrics.

The ability of the MDN to recover these distributional relationships depends strongly on how metabolite information is represented within the model. The use of DreaMS to produce embeddings is central to our approach, as it provides a learned representation of MS/MS spectra that can directly be used in predictive models. DreaMS is one of the available spectral representation-learning methods aimed at encoding mass spectra into latent spaces capturing chemical similarity. Related approaches include Spec2Vec (Huber, Ridder, et al., 2021) and MS2DeepScore (Huber, Burg, et al., 2021), which similarly learn spectral embeddings but differ in their training objectives and architectures. A strength of DreaMS is its self-supervised pretraining on large-scale spectral datasets, allowing it to capture chemical information beyond the molecular coverage of annotated spectral libraries typically used by similarity-based methods (Bushuiev et al., 2026). However, a limitation is that the quality of the embeddings depends on the diversity and coverage of the spectra used during pretraining.

Although DreaMS embeddings provide an effective representation for this task, alternative approaches for generating metabolite representations are also possible. One alternative is to first infer molecules and then generate embeddings using those inferences using tools such as SIRIUS 4 (Dührkop et al., 2019). This approach has the appeal of producing chemically interpretable representations grounded in known molecular structure, but it introduces two inference steps that each carry their own uncertainty. Incorrect or ambiguous molecular annotations can propagate into the embeddings and cause misleading downstream associations. Since many features remain unannotated and annotation confidence can vary widely, bypassing structure inference and directly working with spectral embeddings reduces the risk of compounding errors and allows metabolites to contribute to model training regardless of whether they can be identified.

These considerations also raise the question of how embedding representations can be optimized for downstream prediction tasks. Fine-tuning may represent an important direction for improving predictive performance in this system. Different embedding methods capture different aspects of chemical similarity, and their usefulness for prediction tasks can vary substantially depending on model architecture and training objective. In our study, the MDN was trained on fixed DreaMS embeddings. Prior work on chemical language models has shown that frozen embeddings often perform substantially worse than full-model fine-tuning, suggesting that pretrained representations encode complex nonlinear relationships that are difficult to extract without adaptation (Feller et al., 2026).

Beyond the choice of spectral representation, mixture density networks themselves introduce several important modeling decisions, particularly regarding the number of Gaussian mixture components. Increasing the number of components generally increases model flexibility and allows the network to represent more complex or multimodal distributions. However, larger numbers of components can also increase the risk of overfitting, producing distributions that are excessively peaked or that begin to capture noise rather than meaningful structure (Supplementary Figure S1). This tradeoff is similar to selecting the number of bins in a histogram: too few bins can oversimplify the underlying structure, while too many bins can emphasize noise and create misleading patterns. As a result, selecting the number of mixture components often requires a degree of human judgment in balancing interpretability, flexibility, and generalization performance. In other applications of MDNs, the Akaike information criterion (AIC) and the Bayesian information criterion (BIC) were used to select the number of components (Li et al., 2024). In our case, we evaluated models with one to five, 10, 15, and 20 components using the rare-features training set and selected 15 components as the configuration that best balanced negative log-likelihood and distributional smoothness (Supplementary Figure S2). This number was also used for the all-features model for consistency. Although validation losses continued to decrease at 20 components, the improvement relative to 15 components was modest and was not considered sufficient to justify the increased risk of overfitting. More principled selection approaches could be explored in future work to reduce reliance on manual tuning.

Some limitations likely arise from characteristics of the dataset itself. Some of the environmental variables within the Panama dataset showed constrained variation. Although many metabolites were associated with multiple environmental measurements, the values span relatively narrow ranges. This limits the information available for the model to distinguish feature-specific environmental associations and can encourage convergence toward shared distributional shapes. Additionally, having data for only 44 plots concentrated within a relatively small geographic region restricts the diversity of environmental contexts that a given metabolite can experience. These constraints likely contributed to the observed similarity among the predicted feature-level distributions and highlight the need for broader spatial sampling in future work.

In addition to limited environmental variation, the discrete sampling of underlying continuous environmental gradients introduced another modeling challenge. Measurements were available only for the 44 forest plots included in the study, resulting in discontinuities between observed values. These gaps do not necessarily represent environmental conditions in which a metabolite is absent, but rather conditions that were not represented by the sampled plots. This discrete sampling produced sharply multimodal observed distributions, a structure that can pose difficulties for MDNs because they assume that the target distribution can be approximated by a mixture of Gaussian components (Bishop, 1994). As a result, the model may fit too closely to the discrete observations rather than generalizing across unsampled portions of the environmental gradient. Consequently, gaps in the predicted distributions may reflect limitations in environmental sampling rather than true ecological boundaries in metabolite occurrence.

Together, these limitations suggest several directions for future work. Applying MDNs to larger datasets spanning broader geographic and environmental gradients could help improve predictive performance and allow for a more robust evaluation of the model’s capabilities. Incorporating additional environmental predictors or alternative spectral representations may further improve the quality of the predicted distributions generated by the model.

Despite the limitations, our findings underscore the value of distributional prediction through mixture density networks in metabolomics. Traditional regression models would collapse multimodal or highly variable relationships into single summary values. In contrast, MDNs preserve information about variability and uncertainty, which are central to understanding how metabolites are distributed across landscapes.

### Data availability

The code used for data processing, model training, and analysis is available on GitHub at https://github.com/kellyktvt/Panama_forest_analysis. Spectra were deposited as a public MassIVE dataset (Dataset ID: MSV000104086) on the GNPS server (https://massive.ucsd.edu). Panama forest plot census data containing the distribution and abundance of each species are available on Dryad (https://doi.org/10.15146/mdpr-pm59), and the soil phosphorus and dry-season moisture deficit data for each plot are available on Smithsonian figshare (https://smithsonian.figshare.com/articles/dataset/_Dataset_Tree_Species_Occurrence_Related_to_Dry_Season_Duration_Panama_Canal_Area/25620852).

## Acknowledgments

This work was supported by a grant from Corteva Agrisciences to the Smithsonian Tropical Research Institute, US National Science Foundation award DEB 2240430, University of Texas at Austin Stengl-Wyer Endowment Grant SWG-23-03, Lee and Linda Steen Norris, Chris Harte, and a Stengl-Wyer Scholars fellowship to DLF. C.O.W. also acknowledges support from the Blumberg Centennial Professorship in Molecular Evolution and from Texas Biologics for the BioML Consulting program.

## Supplementary figures

### Supplementary files

**File S1:** Actual and predicted distributions for dry-season moisture, log resin phosphorus, and log total phosphorus across different number of prediction samples.

File: actual vs predicted sample comparison.pdf

**File S2:** Actual and predicted distributions of 180 random features for dry-season moisture. File: random features DrySeasonDeficit 10000.pdf

**File S3:** Actual and predicted distributions of 180 random features for log resin phosphorus. File: random features ResinPi 10000.pdf

**File S4:** Actual and predicted distributions of 180 random features for log total phosphorus. File: random features TotalP 10000.pdf

**File S5:** Actual and predicted distributions of every rare feature with 4 observations for dry-season moisture.

File: rare features 4 observations DrySeasonDeficit 10000.pdf

**File S6:** Actual and predicted distributions of every rare feature with 4 observations for log resin phosphorus.

File: rare features 4 observations ResinPi 10000.pdf

**File S7:** Actual and predicted distributions of every rare feature with 4 observations for log total phosphorus.

File: rare features 4 observations TotalP 10000.pdf

---

**Figure S1:**
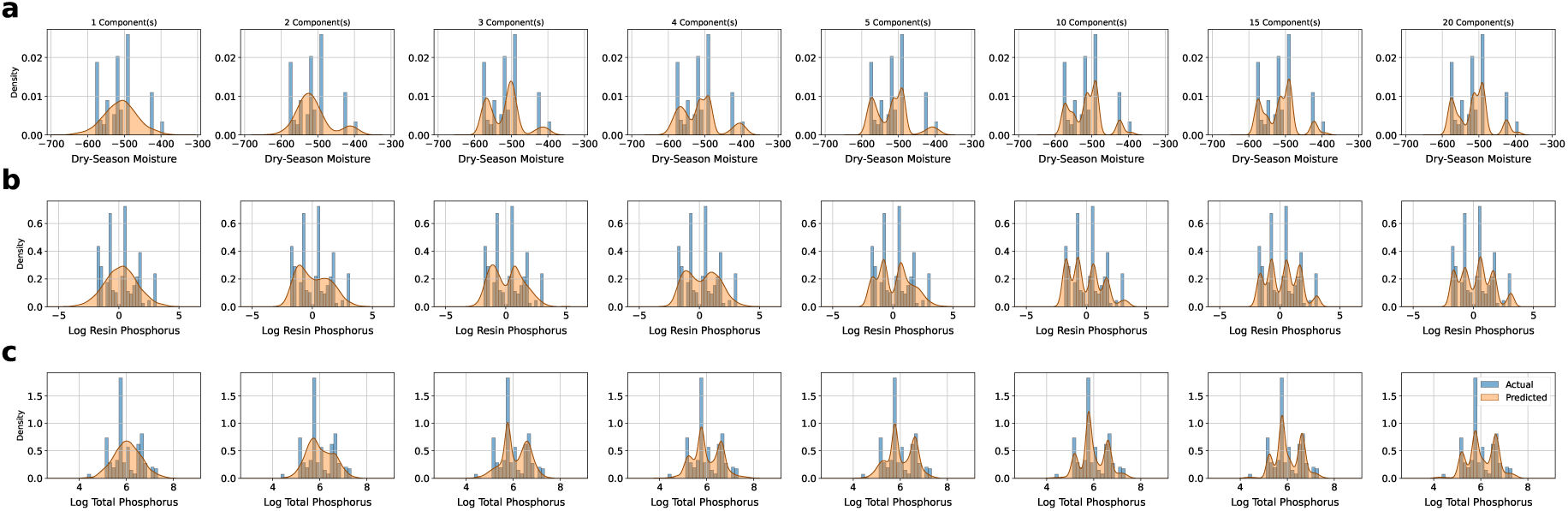
As the number of Gaussian mixture components increases, the peaks of the predicted distributions become more pronounced. The blue bars show the actual values from the test set for each target variable. The orange curves show the distributions of the model’s predictions. Predicted distributions were constructed by generating one prediction per test observation. (a) Actual and predicted distributions for dry-season moisture across different number of components. (b) Actual and predicted distributions for log resin phosphorus across different number of components. (c) Actual and predicted distributions for log total phosphorus across different number of components.

**Figure S2:**
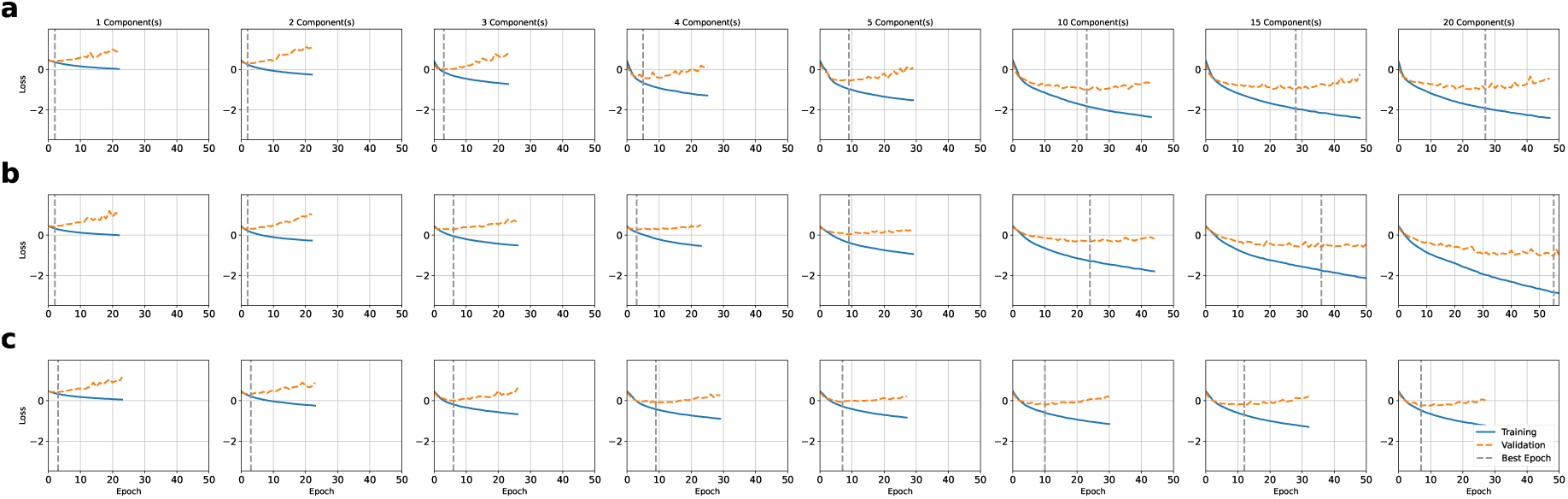
Training and validation loss curves for the 3 target variables after training on rare features using 8 different numbers of components (1-5, 10, 15, 20). The blue solid lines show the training loss, while the orange dashed lines show the validation loss. The gray dashed vertical lines indicate the epochs with the lowest validation loss. (a) Loss curves for dry-season moisture. (b) Loss curves for log resin phosphorus. (c) Loss curves for log total phosphorus.

