## Supplementary figures and images for "Using Mixture Density Networks to Predict Plot-level Environmental Factors from Tree Secondary Metabolites"

### File S1

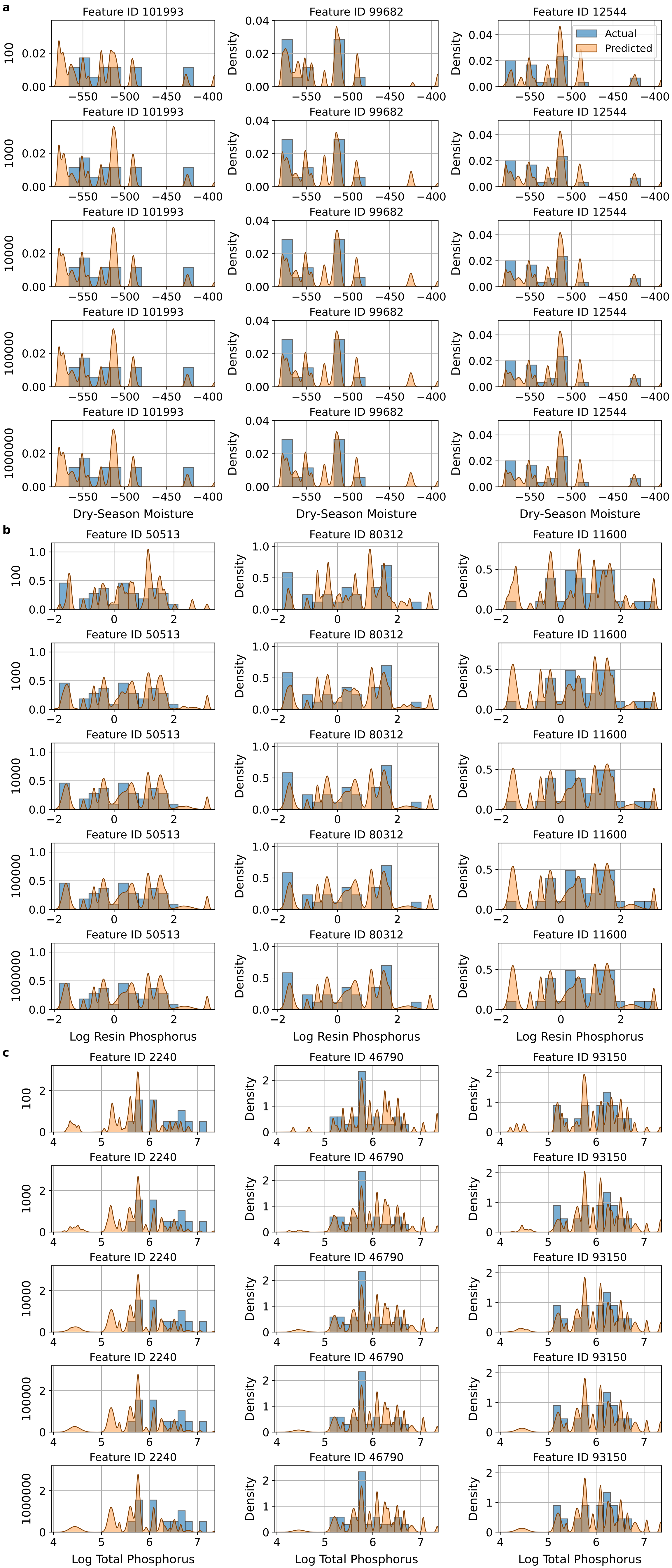

### File S2

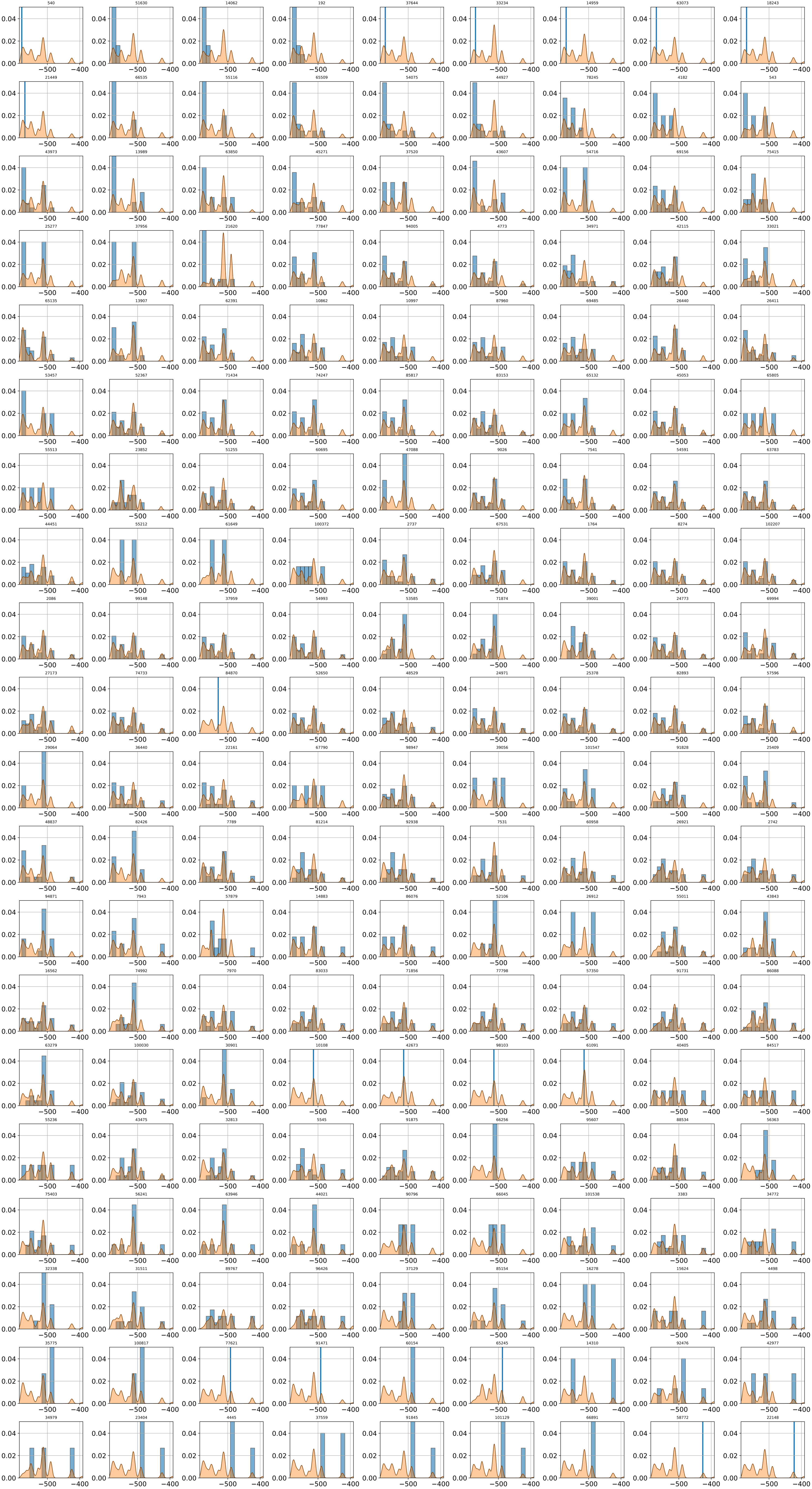

### File S3

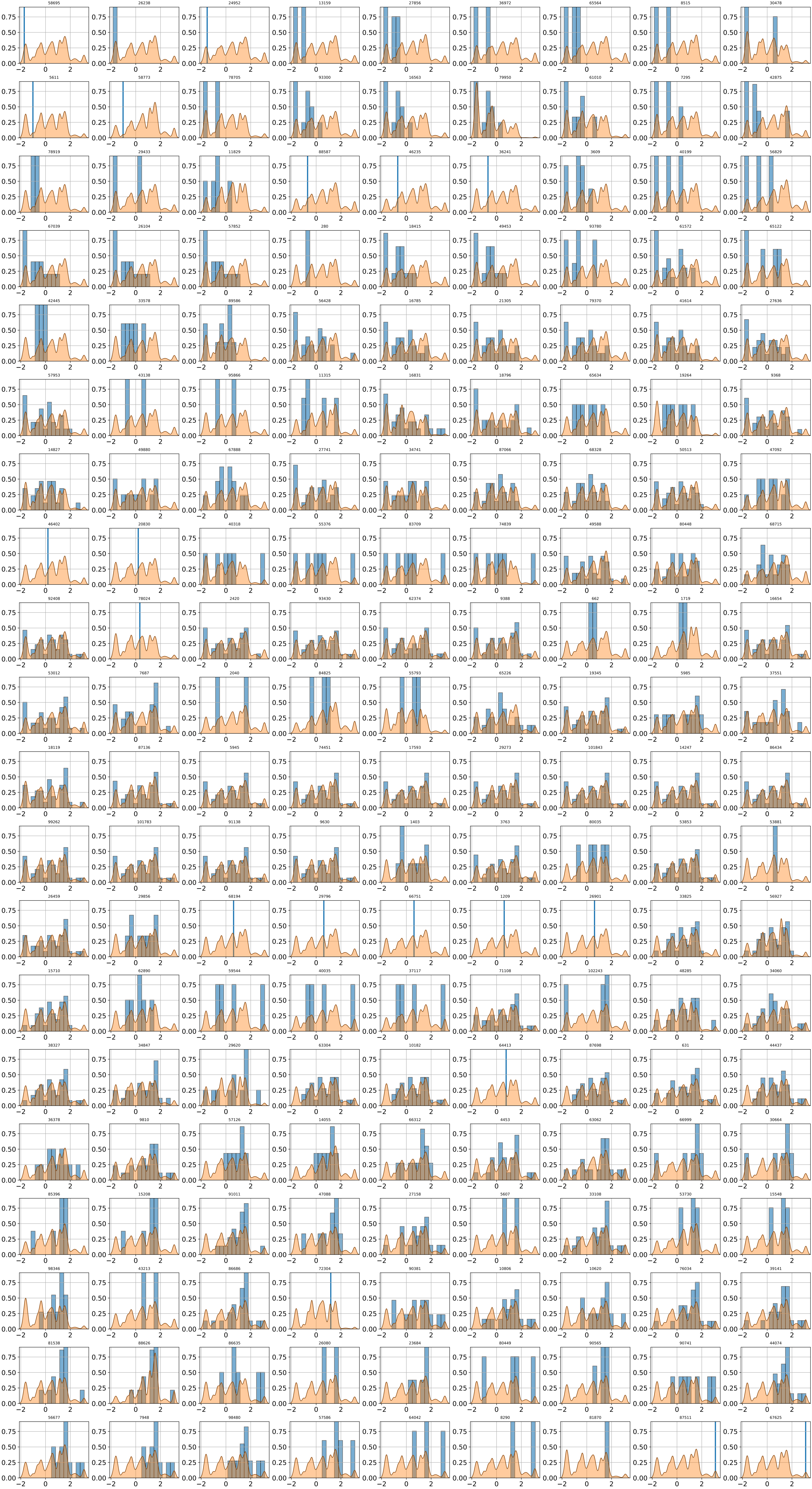

### File S4

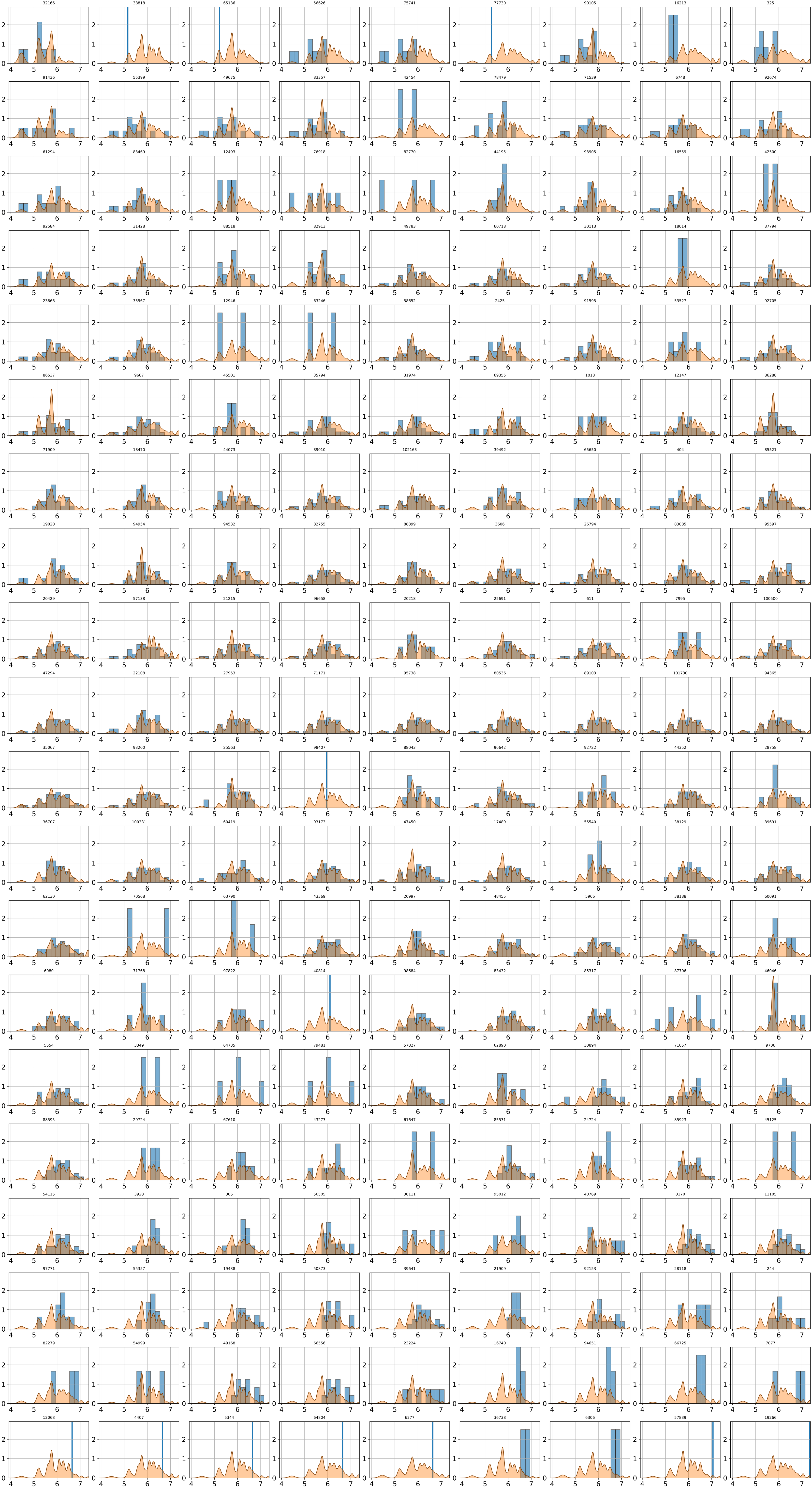

### File S5

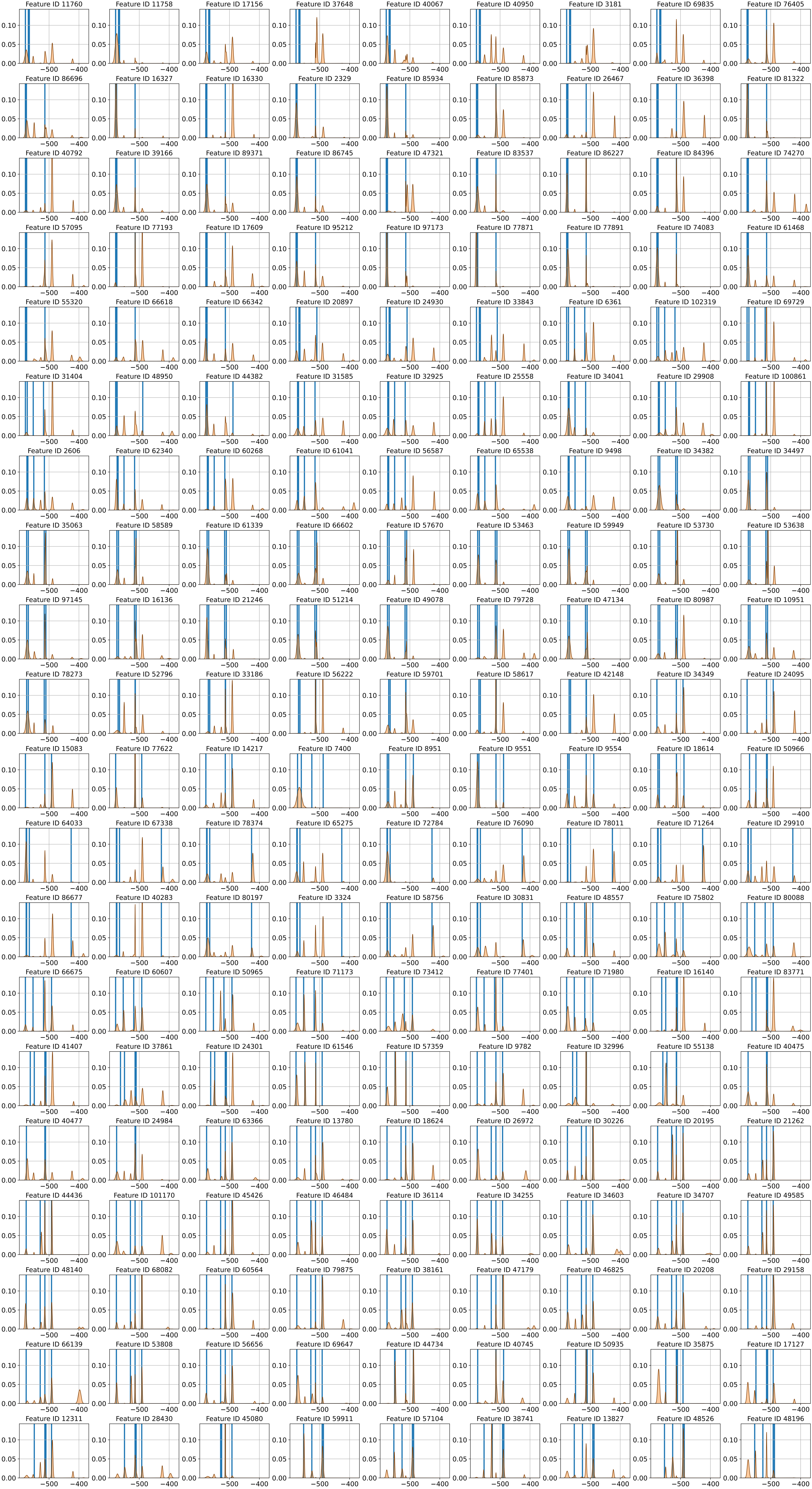

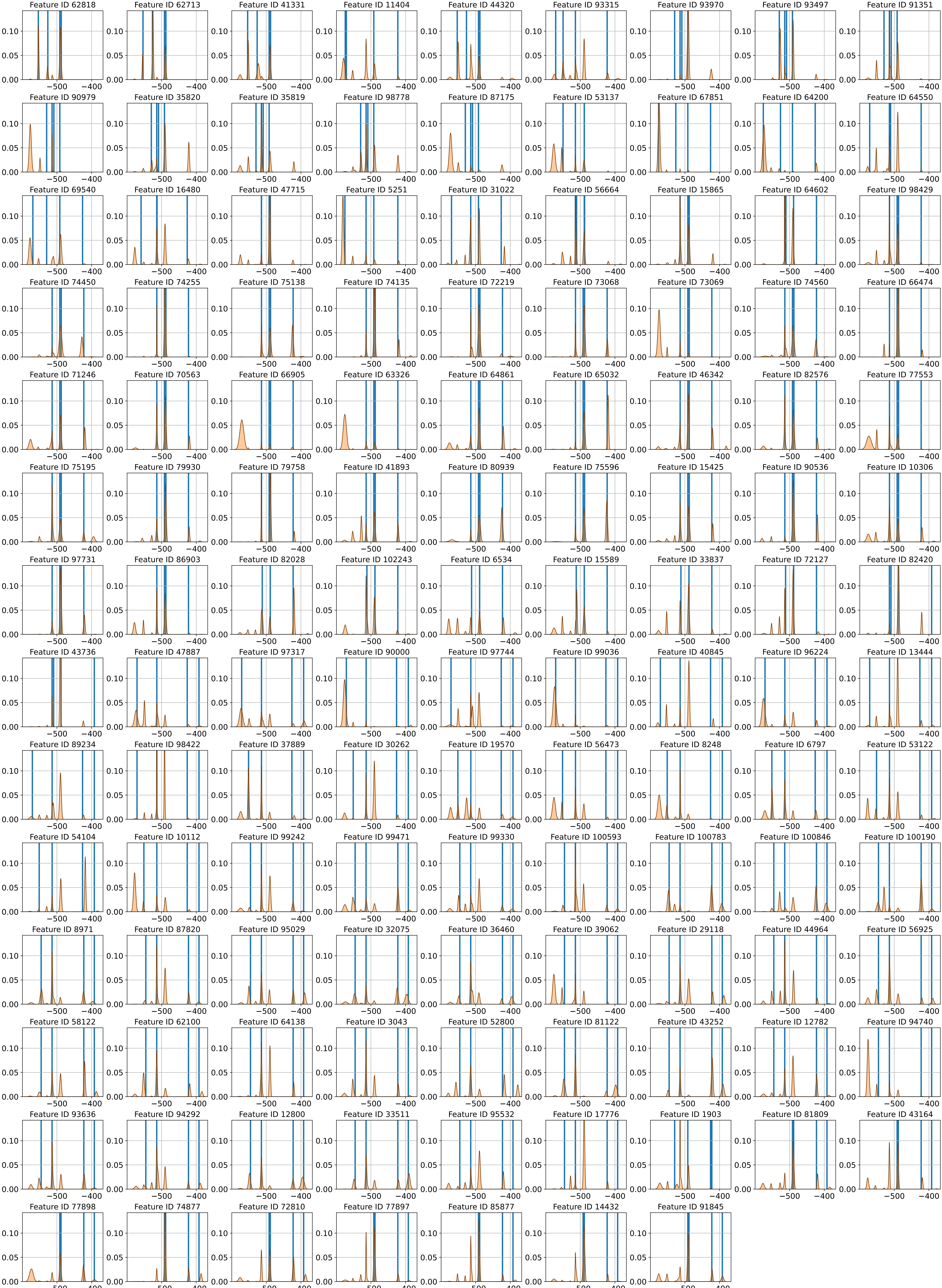

### File S6

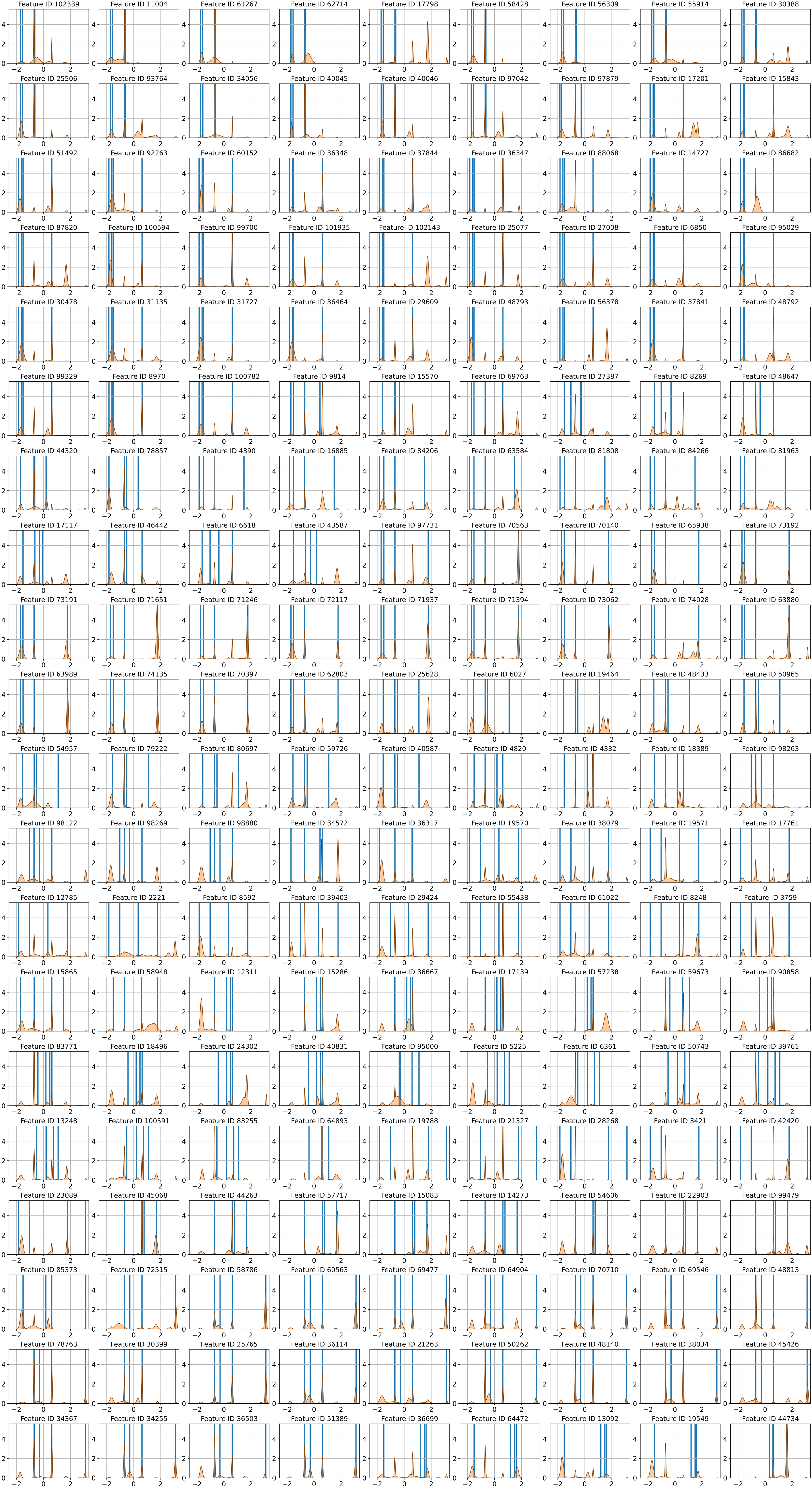

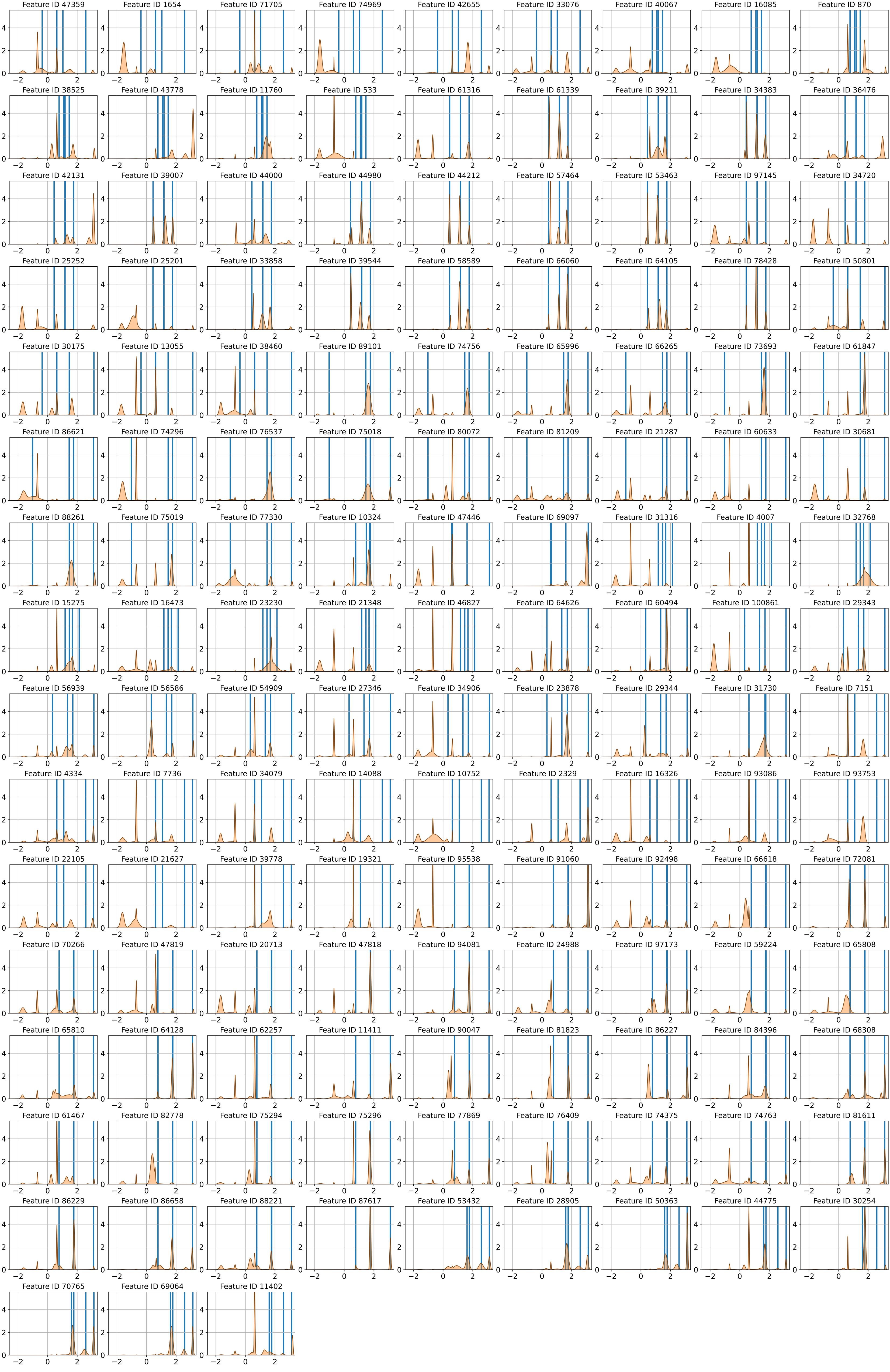

### File S7

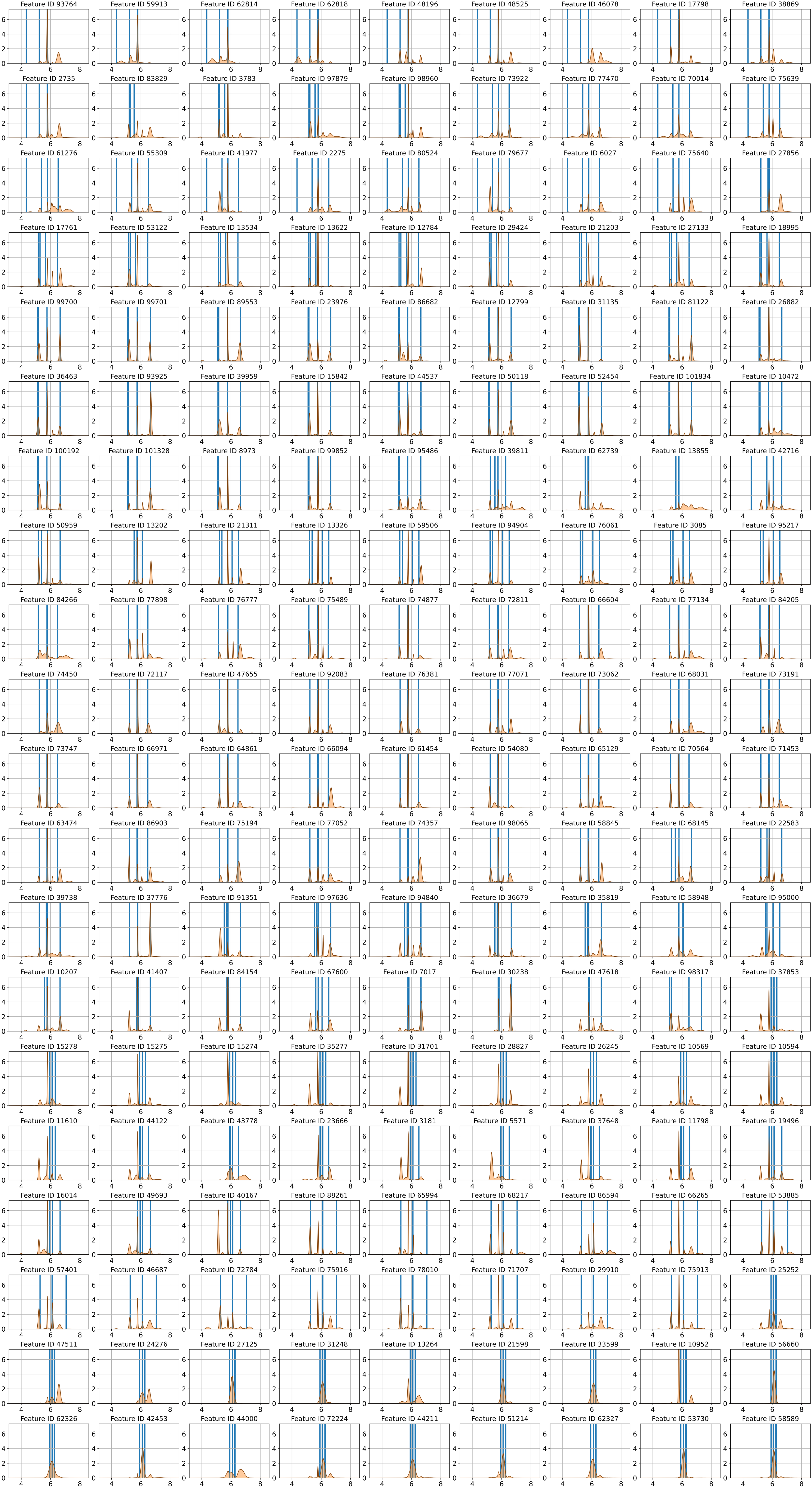

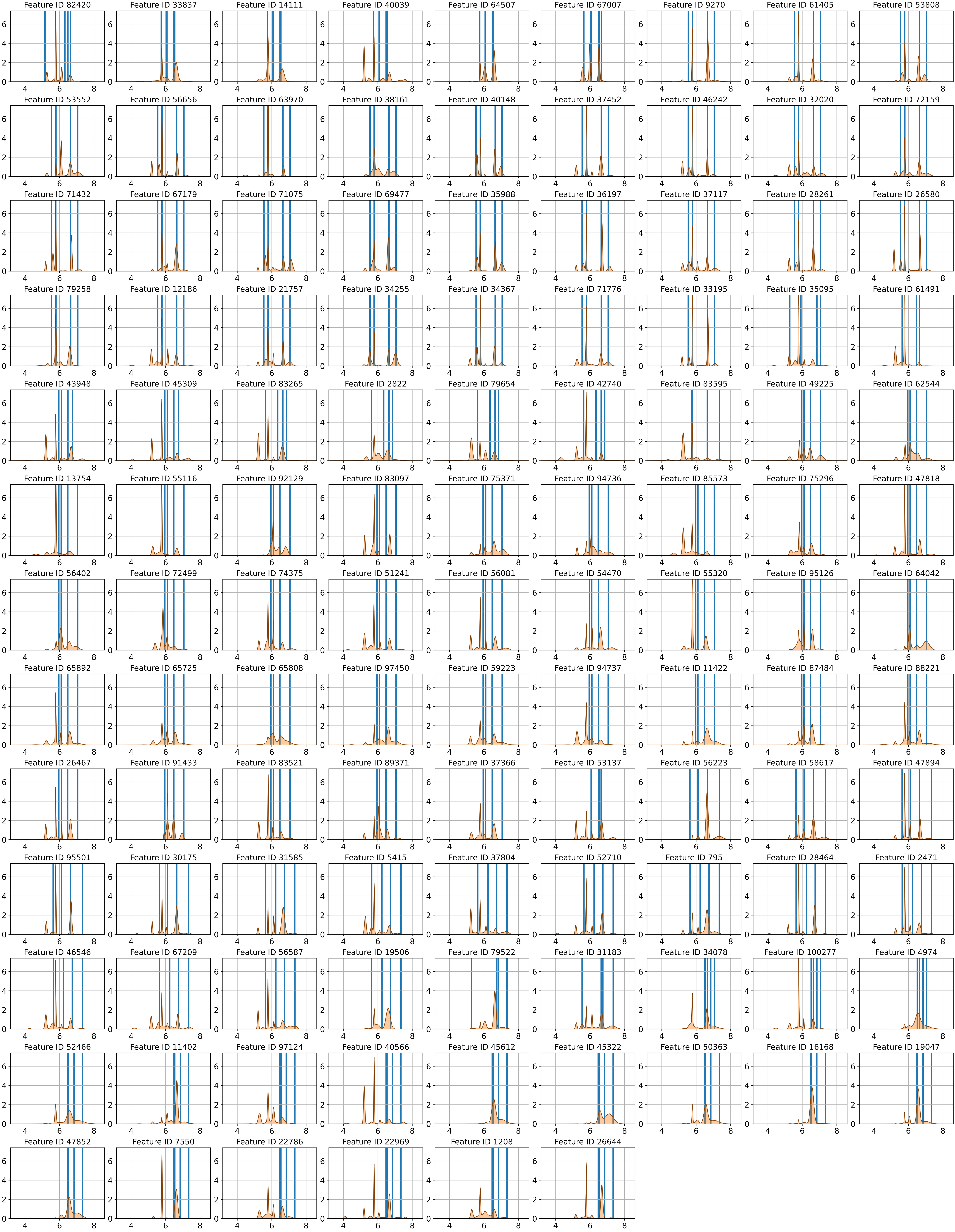
